# Differential regulation of progranulin derived granulin peptides

**DOI:** 10.1101/2021.01.08.425959

**Authors:** Tingting Zhang, Huan Du, Mariela Nunez Santos, Xiaochun Wu, Thomas Reinheckel, Fenghua Hu

## Abstract

Haploinsufficiency of progranulin (PGRN) is a leading cause of frontotemporal lobar degeneration (FTLD). PGRN is comprised of 7.5 granulin repeats and is processed into individual granulin peptides in the lysosome. However, very little is known about the levels and regulations of individual granulin peptides due to the lack of specific antibodies. Here we report the generation and characterization of antibodies specific to each granulin peptide. We found that the levels of granulins C, E and F are differentially regulated compared to granulins A and B. Furthermore, we demonstrated that granulin B, C and E are heavily glycosylated and the glycosylation pattern of granulin C varies in different physiological and pathological conditions. Deficiency of lysosomal proteases leads to alterations in the levels of a specific subset of granulins. These data support that the levels of individual granulin peptides are differentially regulated under physiological and pathological conditions and provide novel insights into how granulin peptides function in the lysosome.

## Introduction

Progranulin (PGRN) protein, encoded by the *granulin (GRN)* gene, has been implicated in several neurodegenerative diseases (Bateman & Bennett, 2009; Cenik *et al*, 2012). Haplo-insufficiency of the protein, due to heterozygous mutations in the *GRN* gene, is a leading cause of frontotemporal lobar degeneration with TDP-43 aggregates (FTLD-TDP) (Baker *et al*, 2006; Cruts *et al*, 2006; Gass *et al*, 2006). Homozygous PGRN mutations result in neuronal ceroid lipofuscinosis (NCL), a lysosomal storage disorder (Almeida *et al*, 2016; Smith *et al*, 2012). PGRN is known as a secreted glycoprotein of 7.5 granulin repeats (Bateman & Bennett, 2009; Cenik *et al.*, 2012). However, accumulating evidence has suggested a critical role of PGRN in the lysosome (Paushter *et al*, 2018). PGRN deficiency has been shown to result in lysosome abnormalities with aging (Ahmed *et al*, 2010; Tanaka *et al*, 2014). At the molecular and cellular level, PGRN is a lysosome resident protein (Hu *et al*, 2010) and is transcriptionally co-regulated with many essential lysosomal genes by the transcriptional factor TFEB (Belcastro *et al*, 2011; Sardiello *et al*, 2009). PGRN interacts with another lysosomal protein prosaposin (PSAP) to facilitate each other’s lysosomal trafficking (Hu *et al.*, 2010; Zhou *et al*, 2015; Zhou *et al*, 2017c). Within the lysosome, PGRN has been shown to get processed to granulin peptides by cathepsins (Holler *et al*, 2017; Lee *et al*, 2017; Zhou *et al*, 2017b). These granulin peptides have been proposed to possess unique biological activities, in a way similar to the saposin peptides derived from PSAP, which function as activators for enzymes involved in glycosphingolipid degradation (O’Brien & Kishimoto, 1991). In line with this, PGRN and granulin peptides have been shown to regulate the activities of several lysosome enzymes, including cathepsin D (Beel *et al*, 2017; Butler *et al*, 2019; Valdez *et al*, 2017; Zhou *et al*, 2017a) and glucocerebrosidase (Arrant *et al*, 2019; Valdez *et al*, 2019; Zhou *et al*, 2019).

Despite these studies, very little is known about how granulin peptides are regulated in the lysosome due to the lack of specific antibodies to each individual peptide. In this study, we report the generation and characterization of antibodies to each individual granulin peptide. We show that these antibodies can be used to specifically detect each individual granulin peptide at endogenous levels. Furthermore, we demonstrate that the levels of each granulin peptide vary from one another in physiological and pathological conditions, indicating differences in processing or stability in the lysosome. We also found that granulin B, C and E undergo differential glycosylation. Taken together, these results provide novel insights into the regulation of granulin peptides in the lysosome.

## Results

### Generation of antibodies to each individual granulin peptide

To generate antibodies specific to each granulin peptide, we purified recombinant GST tagged mouse granulin peptides from bacteria (Table 1). These proteins were then used to immunize rabbits to generate polyclonal antibodies. To test the specificity of these antibodies toward each individual granulin peptide, we expressed individual mouse granulin peptides in HEK293T cells with an N-terminal signal sequence followed by a GFP tag. HEK293T lysates containing GFP tagged granulins were then used in western blot analysis to determine the specificity of the antibodies against each individual granulin peptide. All the granulin antibodies specifically recognized their respective granulin peptide, except the granulin B antibody, which exhibits weak cross-reactivities with granulin C (Fig.1A).

**Table 1:**
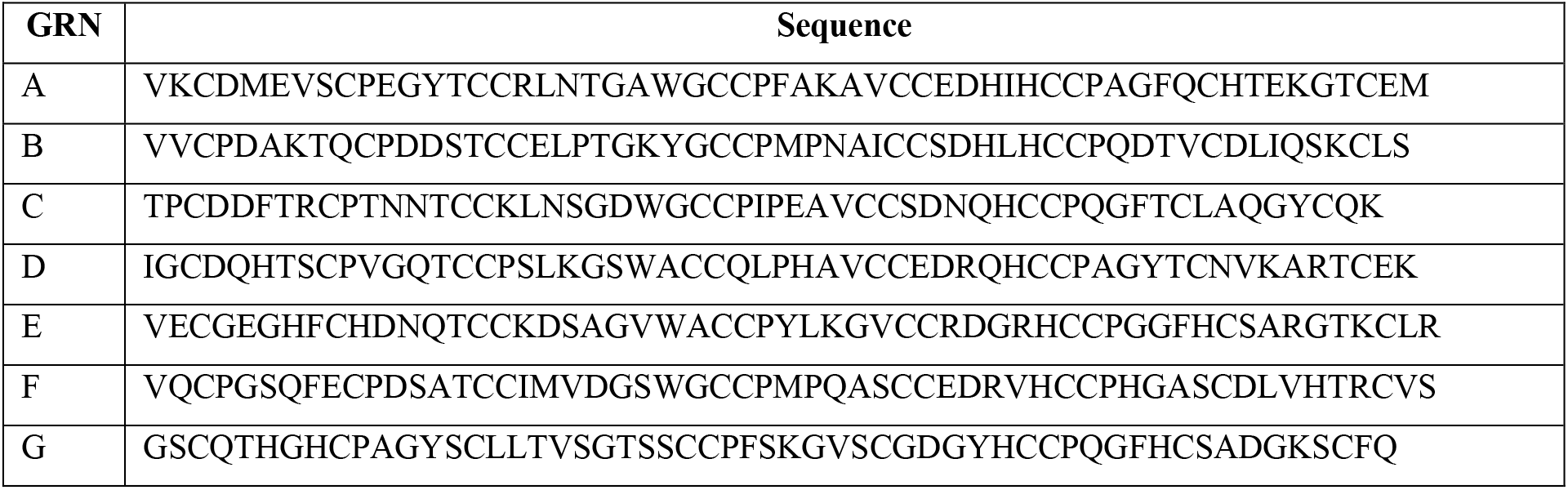
Sequence of mouse granulin peptides used in our study.

**Figure 1.**
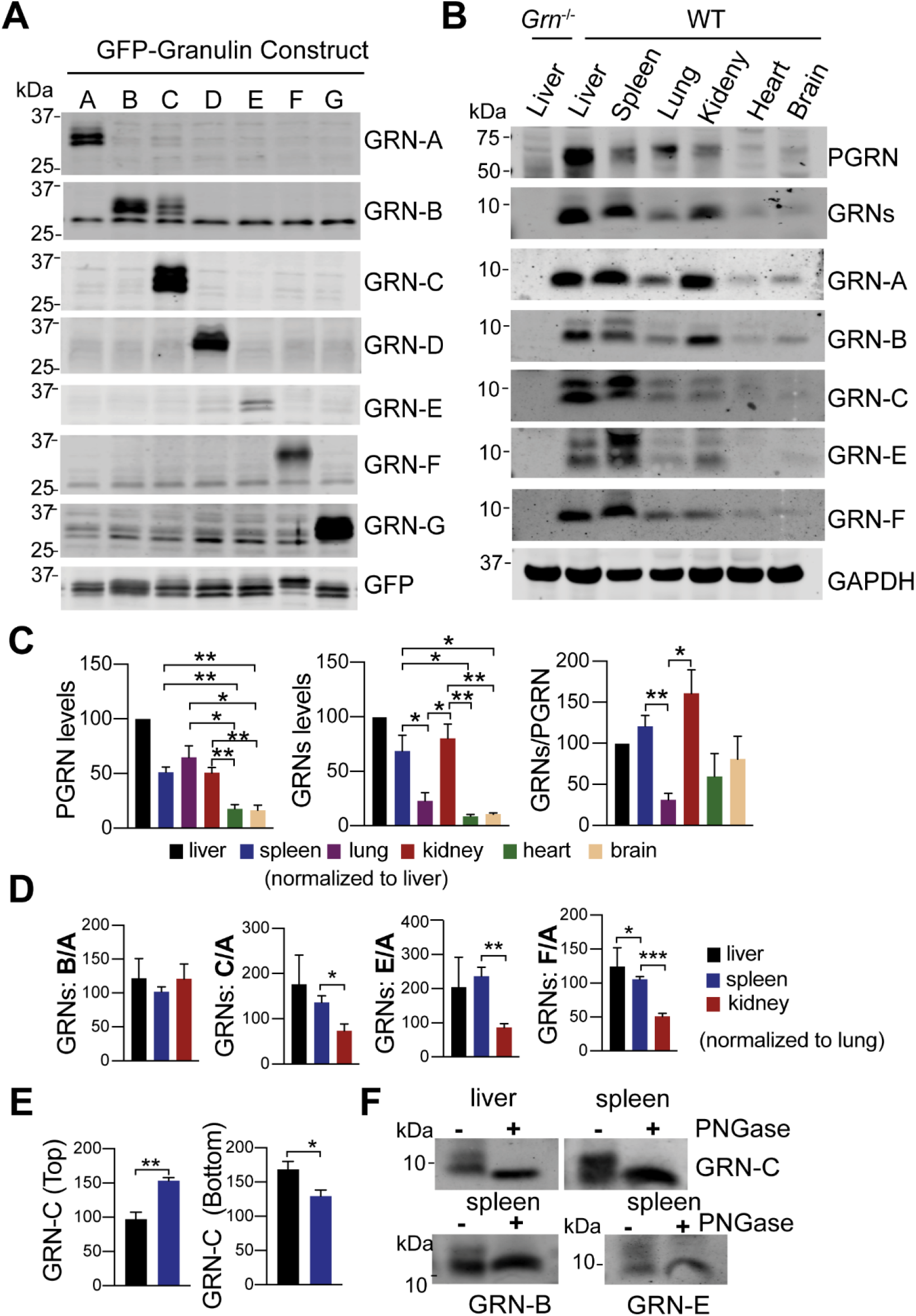
Characterization of granulin antibodies and analysis of granulin levels in tissue lysates. **(A)** HEK293T lysates containing GFP or GFP tagged mouse granulins were probed with antibodies against each individual granulin peptide as indicated. **(B)** Western blot analysis of different tissues lysates from 4-5 month old WT and *Grn*^−/−^ mice with antibodies against each individual granulin peptide as indicated. Mixed male and female mice were used for this analysis. **(C)** Quantification of experiment in (**B**). The ratio between PGRN or granulins and GAPDH as well as between granulins and full length PGRN as recognized by the sheep anti-PGRN antibodies (R&D) was quantified and normalized to that in the liver sample on the same gel (set as 100). Data presented as mean ± SEM. n = 3. *, p<0.05, **, p<0.01, ***, p<0.001, ****, p<0.0001, unpaired two-tailed Student’s *t*-test. **(D)** Quantification of the ratios between granulins B/C/E/F and granulin A. The band intensity of different granulins were measured and normalized to GAPDH. The ratios between granulins B/C/E/F and granulin A were calculated and normalized to the lung sample on the same gel (set as 100). Data presented as mean ± SEM. *, p<0.05, **, p<0.01, ***, p<0.001, unpaired two-tailed Student’s *t*-test. **(E)** Quantification of two differentially glycosylated forms of granulin C for experiment in (B). The ratio between top or bottom bands of granulin C and GAPDH in liver and spleen lysates was quantified and normalized to their levels in the lung. Data presented as mean ± SEM. n = 4. **, p<0.01, unpaired two-tailed Student’s *t*-test. **(F)** Granulin B, C and E are glycosylated. Spleen or liver lysates from WT mice were immunoprecipitated using anti-granulin B, C or E antibodies and the immunoprecipitates were treated with PNGase F.

Next, we determined whether these antibodies could detect endogenous granulin peptide using liver lysates from adult WT and *Grn*^−/−^ mice. Specific signals around 10kDa were successfully detected in the WT lysates but not in the *Grn*^−/−^ samples with granulin A, B, C, E, and F antibodies (Fig. 1B). Unfortunately, granulin D and G antibodies cannot detect endogenous granulin peptides, although they recognize overexpressed granulin peptides efficiently. Thus, we focused our effort on granulins A, B, C, E and F for the current study.

### Variation in the levels of granulin peptides in different tissues

To determine whether the levels of granulin peptides vary from each other, first we analyzed the levels of each granulin peptide in different tissues using western blots (Fig. 1B). We found that PGRN is highly expressed in the liver, spleen, lung and kidney (Fig. 1B, 1C). Using the commercial PGRN antibody which preferentially recognizes granulins B, C and F (Fig. S1), a corresponding enrichment of granulins is detected in the liver, spleen and kidney, but not in the lung (Fig. 1B, 1C). Using antibodies against individual granulins, relatively high levels of granulins A and B in liver, spleen and kidney but not in the lung were also observed (Fig. 1B, Fig. S2), indicating that PGRN processing or the stability of granulins A and B is different in the lung versus spleen and kidney. Interestingly, while the levels of granulins C, E and F are also high in liver and spleen and low in the lung, their levels are relatively low in the kidney, as shown by a significant decrease in the ratio between granulins C/E/F and granulin A in the kidney compared to that in the liver and spleen (Fig. 1B and 1D). This suggests that the levels of granulin peptides could differ from each other although they are derived from the same precursor. This could be due to differential processing or differences in their stability within the lysosome.

### Glycosylation of granulins B, C and E

In our western blot analysis, two distinct bands have been observed for granulin B, C and E at endogenous levels (Fig. 1B). More interestingly, the pattern of glycosylation for granulin B and C differs in the spleen lysates versus liver lysates (Fig. 1B,1E). In both cases, an increased level of the highly glycosylated form was observed in the spleen compared to the liver, especially for granulin C (Fig. 1B, 1E). PGRN is predicted to contain 5 N-glycosylation sites with granulin B, C and E each harboring one glycosylation site. Additionally, glycosylation sites in granulins C and E have been mapped by mass spectrometry analysis (Songsrirote *et al*, 2010). We speculated that these two bands observed for granulins B, C and E could be peptides with different degrees of glycosylation. To test that, we immunoprecipitated granulin B, C and E peptides with their corresponding antibodies and treated the immunoprecipitates with PNGase F to remove N-glycans. The two bands collapsed to a single band with lower molecular weight with PNGase F treatment, confirming that granulin B, C and E have two different glycosylated forms (Fig. 1F).

### The levels of PGRN and granulin peptides in different brain regions

PGRN is expressed by both neurons and microglia and broadly distributed in different brain regions and the spinal cord (Matsuwaki *et al*, 2011; Petkau *et al*, 2010). To determine the distribution of granulin peptides in different brain regions, we compared the levels of PGRN and granulins in male and female mouse brain regions using western blot analysis (Fig. 2A). In male mice, the levels of full length PGRN is highest in in the hippocampus, corpus callosum and thalamus, as compared to other regions (Fig. 2A and 2B). The levels of granulins detected with commercial anti-PGRN antibodies and the levels of granulin A and C are also high in the hippocampus and corpus callosum lysates, but relatively lower in the thalamus. In addition, although the level of full length PGRN is low in the cortex, the levels of granulins A and C are highest in the cortex (Fig. 2A and 2B). These results indicate that the levels of both PGRN and granulin peptides vary in different brain regions. The efficiency of PGRN processing or the stability of granulin peptides might be subject to region specific regulations in the mouse brain. However, we were unable to obtain consistent results from female mice. In females, both full length PGRN and the ratio between individual granulin and PGRN show great variability from mouse to mouse (Fig. S3). PGRN gene expression is known to be regulated by estrogen (Chiba *et al*, 2007; Suzuki *et al*, 2009; Suzuki & Nishiahara, 2002; Suzuki *et al*, 1998), which might explain the variability seen in individual females due to changes in estrogen levels.

**Figure 2.**
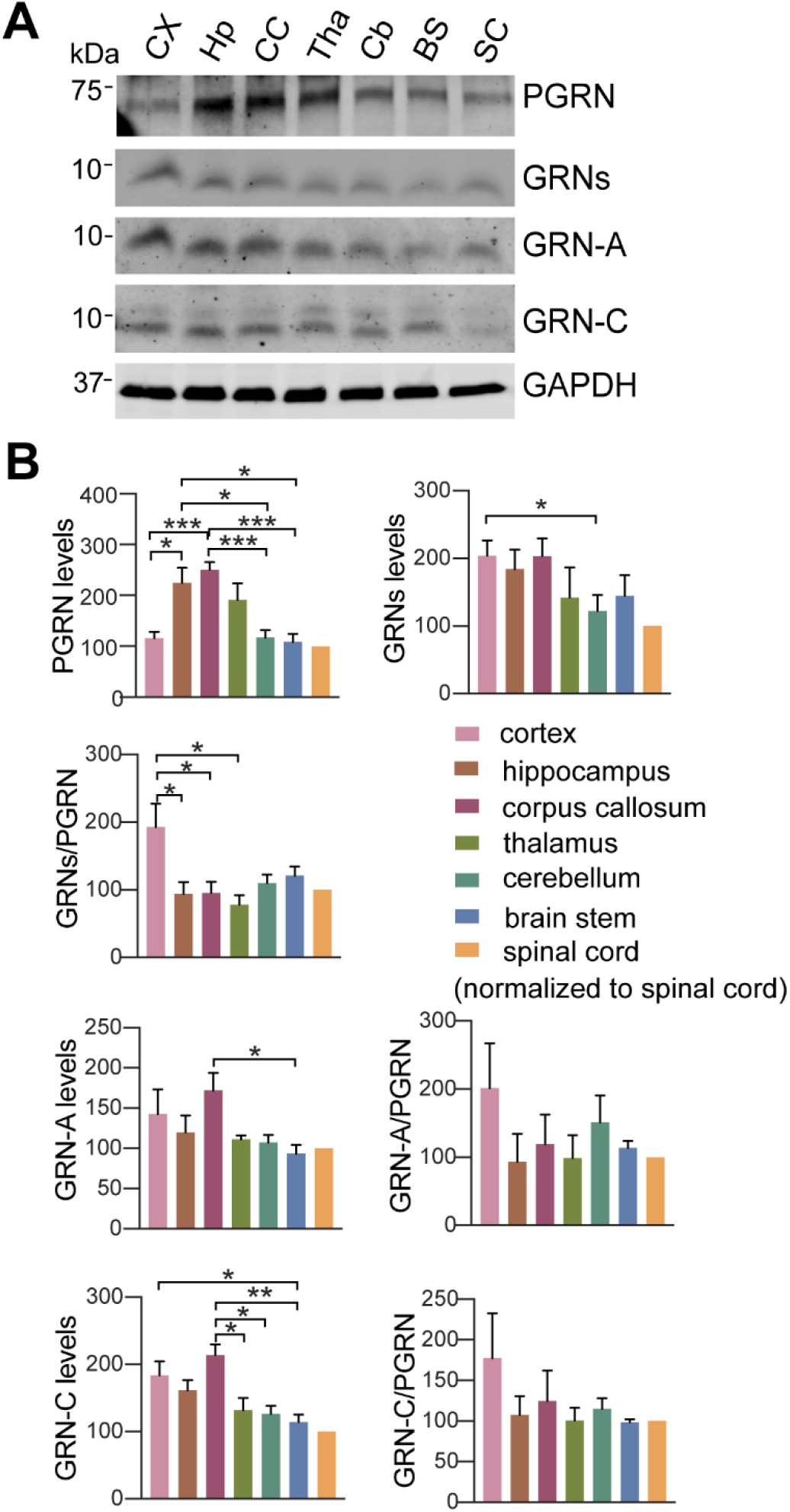
Analysis of PGRN and granulin levels in brain regions and spinal cord. **(A)** Western blot analysis of tissue lysates from 4.5 to 5 month old male WT mice with antibodies against full length of PGRN and individual granulin A and C as indicated. CX = cortex, Hp = hippocampus, CC = corpus callosum, Tha = thalamus, Cb = cerebellum, BS = brain stem, SC = spinal cord. **(B)** Quantification of experiment in (A). The ratio between each granulin peptide to full length PGRN was quantified and normalized to the value of spinal cord samples on the same gel (set as 100). n = 3-5. Data presented as mean ± SEM. *, p<0.05, **, p<0.01, ***, p<0.001, unpaired two-tailed Student’s *t*-test.

### Regulation of PGRN processing and granulin levels by lysosomal proteases

Lysosomal proteases, such as cathepsins, have been shown to play a role in PGRN processing. The cysteine protease, cathepsin L, was shown to cleave PGRN to granulin peptides efficiently in vitro (Holler *et al.*, 2017; Lee *et al.*, 2017; Zhou *et al.*, 2017b). However, how lysosomal proteases regulate PGRN cleavage in vivo remains unclear. Using the granulin specific antibodies, we analyzed the levels of each granulin peptide in the cortex of mice deficient in individual lysosomal proteases, including cathepsin B, D, L, K and Z (Fig. 3A, 3B, 3C). Ablation of most of these cathepsins individually do not seem to have significant effect on the levels of full length PGRN and total granulins, except cathepsin B and cathepsin D. The levels of both PGRN and granulin peptides are significantly upregulated in *Ctsd*^−/−^ cortical lysates (Fig. 3B), partly due to transcriptional up-regulation as reported previously (Gotzl *et al*, 2014). The ratio of granulins A, B, C, but not granulin F, versus full length of PGRN is decreased in the *Ctsd*^−/−^ lysates, suggesting that ablation of cathepsin D affects PGRN processing or the stability of a subset of granulins (Fig. 3C). Interestingly, a significant increase in the levels of heavily glycosylated form of granulin C was observed in *Ctsd*^−/−^ lysates (Fig. 3C), suggesting possible dysfunction of lysosomal glycosidases in response to cathepsin D loss. Although cathepsin B deficiency does not have any obvious effect on the levels of PGRN, it leads to a significant increase in the levels of granulin A and B without any obvious effects on granulin C and F (Fig. 3A), suggesting that cathepsin B might specifically regulate the generation or stability of granulin A and B in the lysosome. Despite results from in vitro studies supporting an important role of cathepsin L in PGRN processing (Holler *et al.*, 2017; Lee *et al.*, 2017; Zhou *et al.*, 2017b), ablation of cathepsin L does not have any obvious effect on the levels of PGRN and granulin peptides in the brain (Fig. 3C). Cathepsin B and L are known to have overlapping functions. In cortical lysates from cathepsin B and L double knockout (*Ctsb^−/−^ Ctsl*^−/−^) mice, levels of full length PGRN are significantly increased (Fig. 3D), possibly due to transcriptional upregulation caused by severe lysosomal abnormalities in these mice (Felbor *et al*, 2002; Sevenich *et al*, 2006). However, the ratio between individual granulin peptides to full length PGRN is not altered in *Ctsb*^−/−^ *Ctsl*^−/−^ brain lysates, indicating that none of these two cysteine proteases is essential for PGRN processing in vivo. It should be noted that the levels of other lysosomal proteases are likely to be changed upon the loss of one or more proteases (Martinez-Fabregas *et al*, 2018). Thus, it is possible that other proteases get upregulated to process PGRN in the absence of cathepsins B and L. Heavily glycosylated form of granulin C accumulates in *Ctsb^−/−^ Ctsl*^−/−^ cortical lysates similar to that in *Ctsd*^−/−^ mice, indicating changes in the activities of lysosomal glycosidases upon lysosomal dysfunction. Interestingly, the glycosylation pattern of granulin B does not appear to be affected in *Ctsb*^−/−^ *Ctsl*^−/−^ cortical lysates, suggesting the glycosylation of granulin B and C are subject to different regulations.

**Figure 3.**
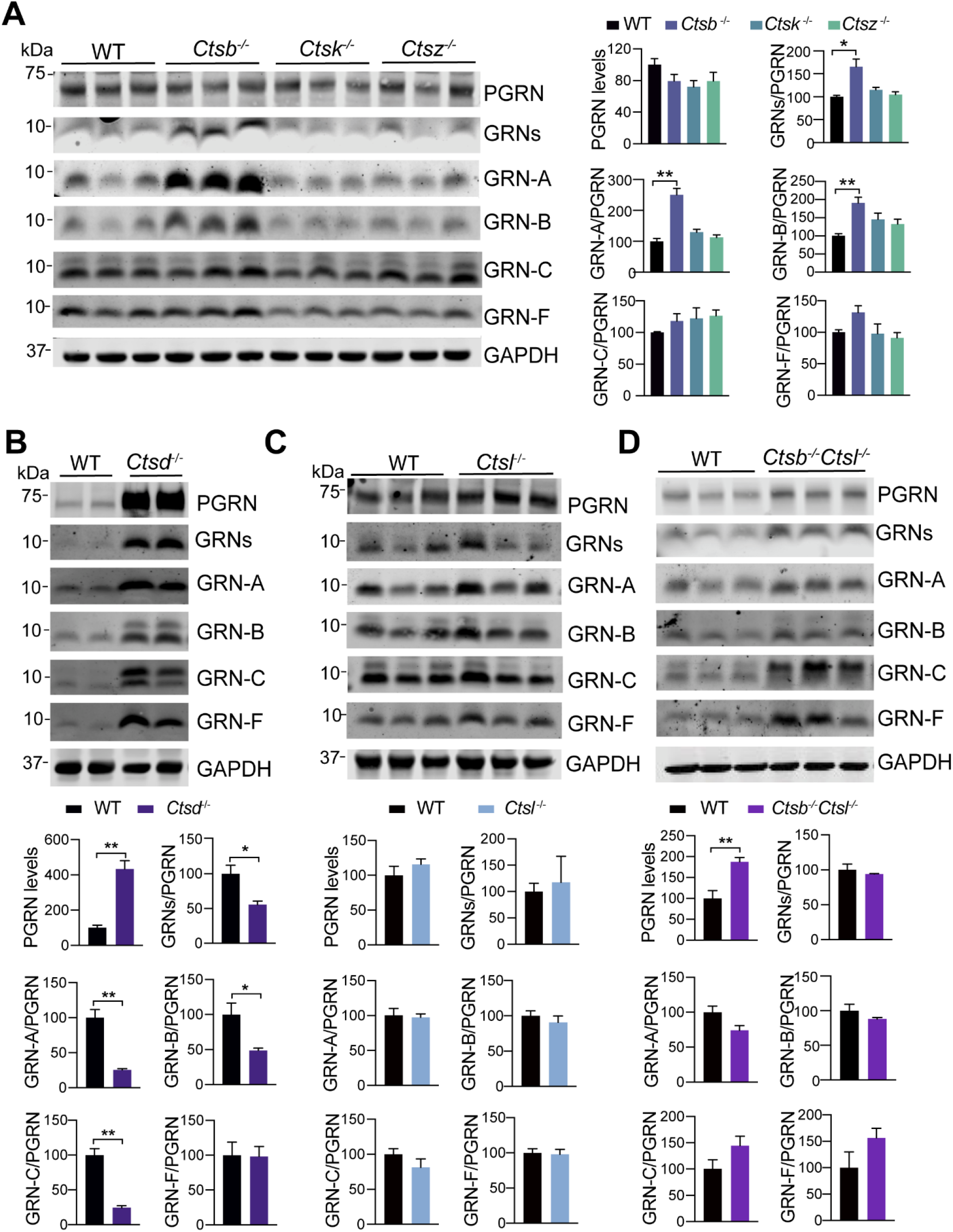
Analysis of PGRN and granulin levels in brain lysates from cathepsin deficient mice. **(A)** Western blot analysis of cortical lysates from 5 month old WT, *Ctsb*^−/−^, *Ctsk*^−/−^ and *Ctsz*^−/−^ mice with antibodies against each individual granulin peptide as indicated. The ratio between each granulin peptide to full length PGRN was quantified and normalized to WT. Data presented as mean ± SEM. n = 3. Data were analyzed by one-way ANOVA tests with Bonferroni’s multiple comparisons. *, p<0.05, **, p<0.01. Mixed male and female mice were used for this analysis. (**B-D**) Western blot analysis of cortical lysates form 2 weeks old *Ctsd*^−/−^ mice (B), 5 month old *Ctsl*^−/−^ mice (C), 3 weeks old *Ctsb*^−/−^*Ctsl*^−/−^ mice (D), and age matched WT mice with antibodies against each individual granulin peptide as indicated. The ratio between PGRN and GAPDH as well as each granulin peptide to full length PGRN was quantified and normalized to WT. Data presented as mean ± SEM. n = 3-5. Data were analyzed by unpaired two-tailed Student’s *t*-test. *, p<0.05, **, p<0.01. Mixed male and female mice were used for this analysis.

## Discussion

PGRN shares many similarities with its binding partner and travel companion prosaposin (PSAP). By forming a complex, these two proteins facilitate each other’s lysosomal trafficking (Hu *et al.*, 2010; Zhou *et al.*, 2015; Zhou *et al.*, 2017c). In addition, when reaching the lysosome, PGRN and PSAP are processed into granulins and saposins, respectively, through the action of lysosomal proteases (Paushter *et al.*, 2018). Saposins are key regulators of enzymes involved in glycosphingolipid degradation pathway (O’Brien & Kishimoto, 1991). Interestingly, although derived from the same precursor, individual saposins are known to be regulated differently in the lysosome. For example, saposin A and D are the main protein components of lipofuscin found in many lysosomal storage diseases (Tyynela *et al*, 1993), indicating that saposin A and D have distinct biochemical properties in the lysosome compared to saposin B and C. However, not much is known about the function and regulation of granulin peptides in the lysosome. Since granulin peptides are likely to be the functional units of PGRN within the lysosome and PGRN haploinsufficiency in FTLD is known to cause haploinsufficiency of granulin peptides (Holler *et al.*, 2017), it is critical for us to understand how the levels of individual granulins are regulated. In this manuscript, we report the generation and characterization of antibodies towards each individual mouse granulin. With these unique antibodies, we have shown that (1) The levels of granulins A and B are differently regulated compared to the levels of granulins C, E and F; (2) Granulins B, C and E are heavily glycosylated and the glycosylation pattern is subject to regulation. Due to the redundancy and cross-regulation of lysosomal proteases, it is challenging to dissect the precise mechanisms involved in PGRN processing. Nevertheless, our results show that cathepsin B might play a role in regulating the levels of granulins A and B and ablation of cathepsin D leads to decreased ratios of granulins A, B and C to PGRN. The generation of antibodies specific to individual granulin peptides allows future work to examine the regulation and function of granulins in physiological and pathological conditions.

## Material and Methods

### Primary Antibodies and Reagents

The following antibodies were used in this study: mouse anti-GAPDH (Proteintech Group, 60004-1-Ig), sheep anti-mouse PGRN (R&D Systems, AF2557) and mouse anti-GFP (Proteintech Group). Fluorescently labelled second antibodies were obtained from Li-Cor and Invitrogen.

To generate polyclonal antibodies to granulin peptides, granulin peptides were cloned in the pGEX6P-1 vector using restriction enzymes BamHI and PmeI (Table 1). The expression construct was transformed into the Origami bacterial strain (Novagen) and the expression of recombinant protein was induced with IPTG. The cells were lysed and lysates were incubated with GST beads. Bound proteins were eluted with glutathione. The buffer was exchanged to PBS using the Centricon devices (Millipore). Recombinant proteins were used to immunize rabbits using services provided by Pocono Rabbit Farm and Laboratory (Canadensis, PA).

The following reagents were also used in the study: Odyssey blocking buffer (LI-COR Biosciences, 927-40000), protease inhibitor (Roche, 05056489001) and Pierce BCA Protein Assay Kit (Thermo scientific, 23225).

### Cell culture

HEK293T were maintained in Dulbecco’s Modified Eagle’s medium (Cellgro) supplemented with 10% fetal bovine serum (Gibco) and 1% Penicillin–Streptomycin (Invitrogen) in a humidified incubator at 37ºC and 5% CO2. Cells were transiently transfected with GFP tagged granulins using polyethylenimine as described (Zhou *et al.*, 2017a). Cells were harvested 2 days after transfection using ice cold RIPA buffer (150 mM NaCl, 50 mM Tris-HCl (pH 8.0), 1% Triton X-100, 0.5% sodium deoxycholate, 0.1% SDS) with 1 mM PMSF, proteinase and phosphatase inhibitors.

### Mouse Strains

C57/BL6 and *Grn*^−/−^ mice (Yin *et al*, 2010) were obtained from The Jackson Laboratory. *Ctsd*−/−(Saftig *et al*, 1995), *Ctsb*−/− (Halangk *et al*, 2000), *Ctsl*−/− (Roth *et al*, 2000), *Ctsb*−/− *Ctsl*−/− (Sevenich *et al.*, 2006), *Ctsk*−/−(Saftig *et al*, 1998) and *Ctsz*−/− (Sevenich *et al*, 2010) mice were characterized previously. All animals (1-6 adult mice per cage) were housed in a 12h light/dark cycle.

### Tissue Preparation and Western blot analysis

Mice were perfused with 1× PBS and tissues were dissected and snap-frozen with liquid nitrogen and kept at −80°C. On the day of the experiment, frozen tissues were thawed and homogenized on ice with bead homogenizer (Moni International) in ice cold RIPA buffer (150 mM NaCl, 50 mM Tris-HCl (pH 8.0), 1% Triton X-100, 0.5% sodium deoxycholate, 0.1% SDS) with 1 mM PMSF, proteinase and phosphatase inhibitors. After centrifugation at 14,000 × g for 15 minutes at 4℃, supernatants were collected. Protein concentrations were determined via BCA assay, then standardized. Equal amounts of protein were mixed with loading buffer with fresh b-mercaptoethanol. Samples were separated by 4-12% Bis-Tris PAGE (Invitrogen) and transferred to 0.2μm nitrocellulose. Western blot analysis was performed as described (Zhou *et al.*, 2015). To quantify the levels of full length PGRN or each granulin peptide (GRN), Image J software was used to measure the density of protein bands (75kDa or 10kDa). These values were normalized to GAPDH. To compare the levels of granulin in different tissues, the levels of granulins were first normalized to GPADH and then normalized to the value of liver, lung or spinal cord samples in the same gel.

### De-glycosylation assay

Spleen and/or liver lysates from WT mice were immunoprecipitated using anti-granulin B, C or E antibodies and the immunoprecipitates were treated with PNGase F (New England Biolabs) according to the manufacturer’s instructions.

### Statistical analysis

All statistical analyses were performed using GraphPad Prism 8. All data are presented as mean ± SEM. Statistical significance was assessed by unpaired two tailed Student’s *t* test (for two groups comparison) or one-way ANOVA tests with Bonferroni’s multiple comparisons (for multiple comparisons). P values less than or equal to 0.05 were considered statistically significant. *, p<0.05, **, p<0.01, ***, p<0.001, ****, p<0.0001.

## Data availability

The data supporting the findings of this study are included in the supplemental material. Additional data are available from the corresponding author on request. No data are deposited in databases.

## Author contributions

T.Z determined changes in granulins in various tissues lysates. X.W. purified all the recombinant GST-granulin proteins for immunization. M.N.S characterized the specificity of granulin antibodies using overexpressed cell lysates. H.D. helped with mouse studies and tissue sample preparation. T. R. provided tissues from cathepsin knockout mice. F.H. supervised the project and wrote the manuscript together with T.Z. All authors have read and edited the manuscript.

## Acknowledgements

We thank Kenton Wu for proofreading the manuscript. This work is supported by NINDS/NIA (R01NS088448 & R01NS095954) and the Bluefield project to cure frontotemporal dementia to F.H.

## Ethical Approval and Consent to Participate

All applicable international, national, and/or institutional guidelines for the care and use of animals were followed. The work under animal protocol 2017-0056 is approved by the Institutional Animal Care and Use Committee at Cornell University.

## Supplemental Material

**Figure S1:**
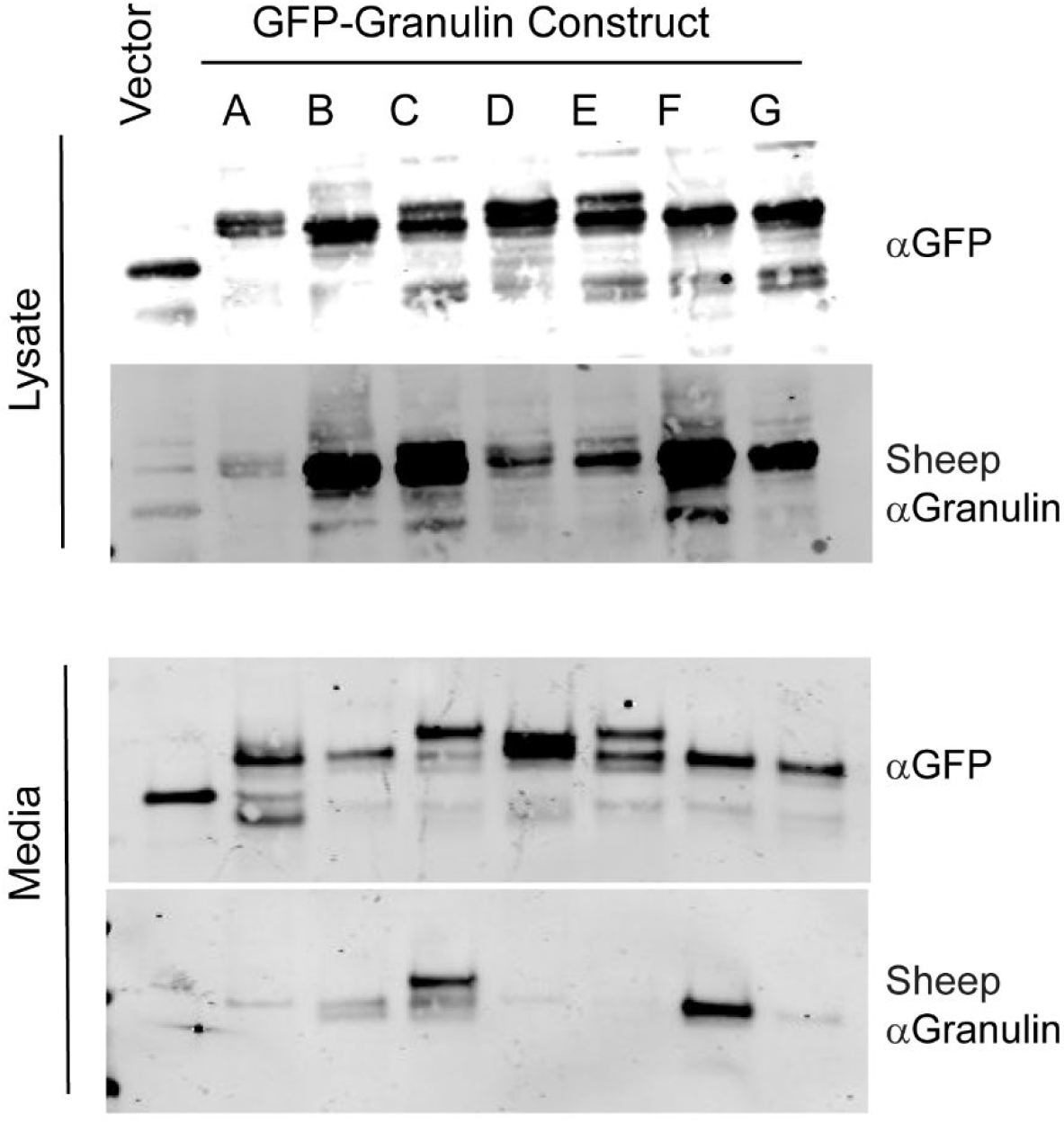
HEK293T lysates and media containing GFP or GFP tagged mouse granulins were probed with sheep anti-mouse PGRN antibodies from R&D systems to detect each individual granulin peptide.

**Figure S2:**
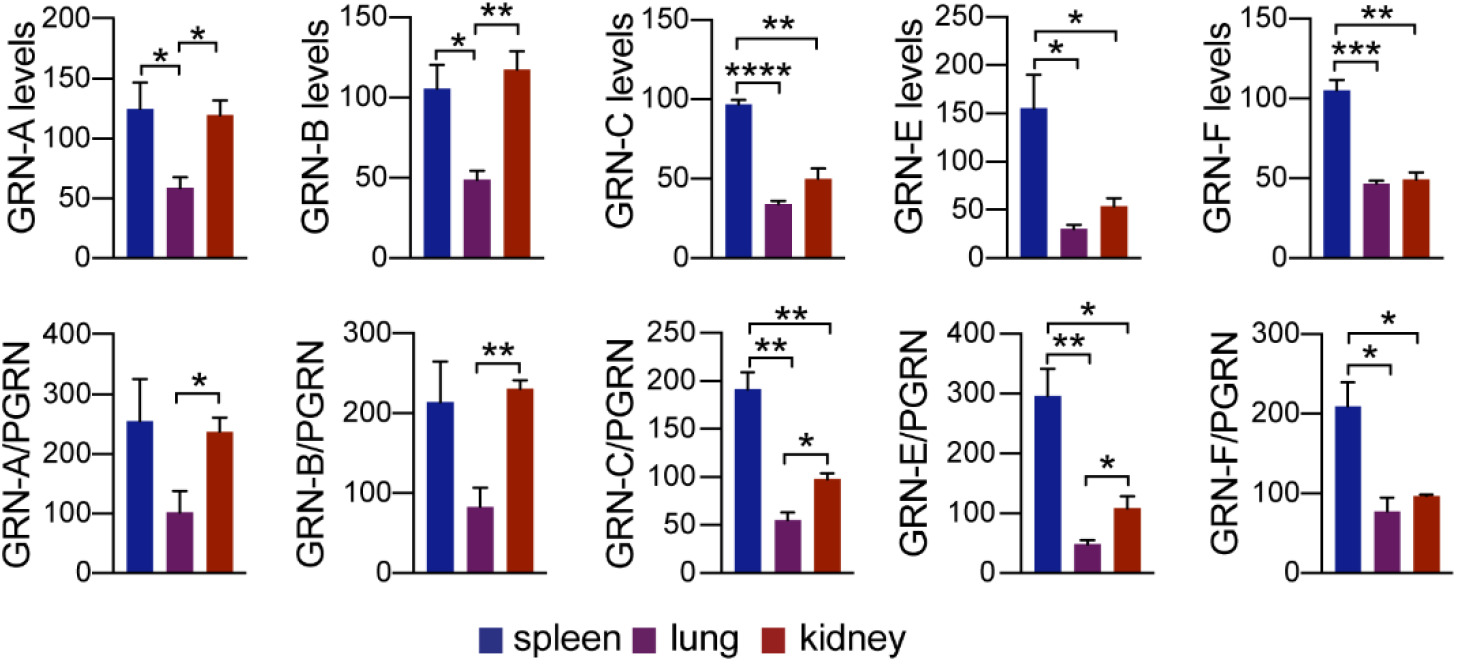
Quantification of the levels of individual granulins and the ratio between granulins and PGRN in the spleen, lung and kidney lysates from 4-5 month old WT and *Grn*^−/−^ mice. The value was normalized to that of liver sample on the same blot (set as 100). Data presented as mean ± SEM. n = 3. *, p<0.05, **, p<0.01, ***, p<0.001, ****, p<0.0001, unpaired two-tailed Student’s *t*-test.

**Figure S3:**
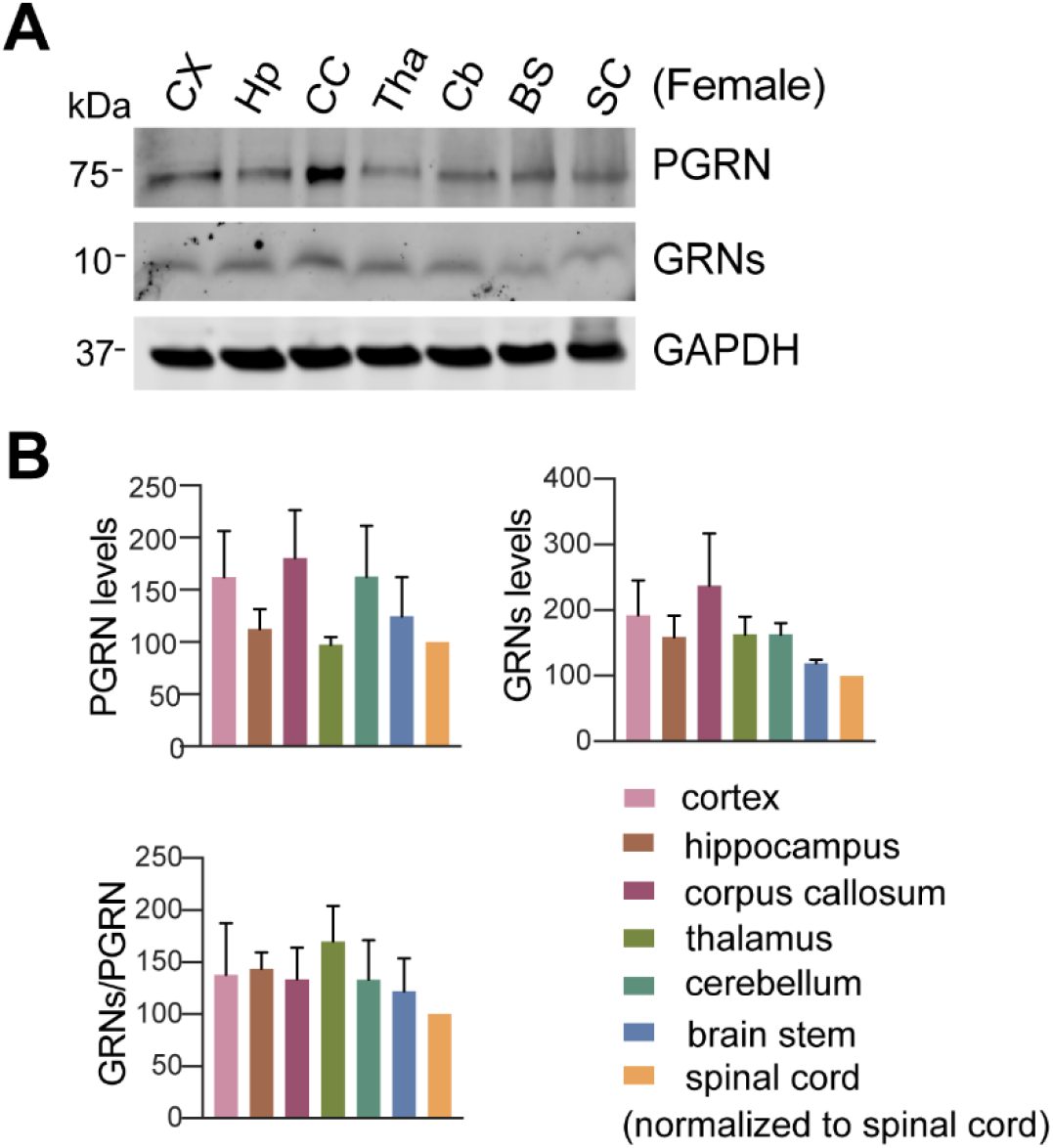
Analysis of PGRN and granulin levels in brain regions and spinal cord of female mice. (**A**) Western blot analysis of tissue lysates from 4.5 to 5 month old female WT mice with antibodies against full length of PGRN and individual granulin A and C as indicated. CX = cortex, Hp = hippocampus, CC = corpus callosum, Tha = thalamus, Cb = cerebellum, BS = brain stem, SC = spinal cord. (**B**) Quantification of experiment in (A). The ratio between total granulins to full length PGRN was quantified and normalized to that of spinal cord. n = 3-4. Data presented as mean ± SEM.

